# SAM-Targeted CRISPR-Cas9 RNP Delivery Combined with Leaf Regeneration Enables DNA-Free, Non-Chimeric Genome Editing in ‘Fuji’ Apple

**DOI:** 10.64898/2026.08.09.742435

**Authors:** Chikako Nishitani, Nozomi Tsujino, Misa Kuroki, Masato Wada, Ryozo Imai

## Abstract

DNA-free genome editing is a promising strategy for the genetic improvement of horticultural crops and fruit trees because it enables targeted mutagenesis without stable genetic transformation. In planta particle bombardment (iPB) delivers CRISPR-Cas9 ribonucleoproteins (RNPs) directly into shoot apical meristems (SAMs), enabling heritable genome editing without the use of tissue culture-based transformation systems. However, the practical application of iPB-mediated editing in fruit trees is limited by the frequent occurrence of chimerism, which cannot be readily eliminated through sexual segregation while maintaining the genetic background of elite cultivars. To overcome this limitation, we combined iPB-mediated RNP delivery with regeneration from edited leaf tissues (iPB-REG). Using this approach, we targeted the self-incompatibility gene S9-RNase in the elite apple cultivar ‘Fuji’ and efficiently recovered non-chimeric edited plants. These results establish iPB-REG as a practical strategy for producing uniform genome-edited fruit trees and provide a valuable platform for DNA-free genetic improvement and functional genomics in clonally propagated perennial crops.

## Introduction

Genome editing has become an important tool for crop improvement because it enables precise modification of target genes. In perennial fruit crops, DNA-free genome editing is particularly attractive because it can generate edited plants without the integration of foreign DNA, potentially facilitating regulatory acceptance and public acceptance.

The in planta particle bombardment (iPB) method enables the direct delivery of CRISPR-Cas9 ribonucleoproteins (RNPs) into shoot apical meristems (SAMs) and has been successfully applied to several plant species (Imai et al., 2020; Kumagai et al., 2022). Despite these advantages, iPB-mediated genome editing frequently generates chimeric plants consisting of edited and non-edited cell lineages. In annual crops, chimerism can often be eliminated by genetic segregation through sexual reproduction. However, this strategy is not suitable for fruit trees because segregation disrupts the highly heterozygous genetic backgrounds that define elite cultivars. Consequently, the development of efficient methods for eliminating chimerism while preserving cultivar identity remains a major challenge for DNA-free genome editing in fruit trees.

Plant regeneration from edited tissues offers a potential solution to this problem by enabling the recovery of plants derived from a limited number of edited cells. In this study, we developed an iPB-regeneration (iPB-REG) strategy to eliminate chimerism after iPB-mediated genome editing in apple and evaluated its utility for recovering non-chimeric edited plants while preserving elite cultivar identity.

## Results and Discussion

The *S-RNase* gene, a key determinant of gametophytic self-incompatibility in apple (Supplementary Fig. 1), was selected as the editing target. Loss-of-function mutations in either *S-RNase* allele confer self-compatibility (Broothaerts et al., 2004; Okada et al., 2024). Two gRNAs were designed to target the *S9-RNase* allele present in ‘Fuji’. Purified recombinant SpCas9 was complexed with chemically synthesized gRNAs to form RNPs, which were subsequently coated onto gold particles and delivered into SAMs prepared from *in vitro*-propagated plantlets by removing newly emerging leaves (Fig. 1a).

**Figure 1.**
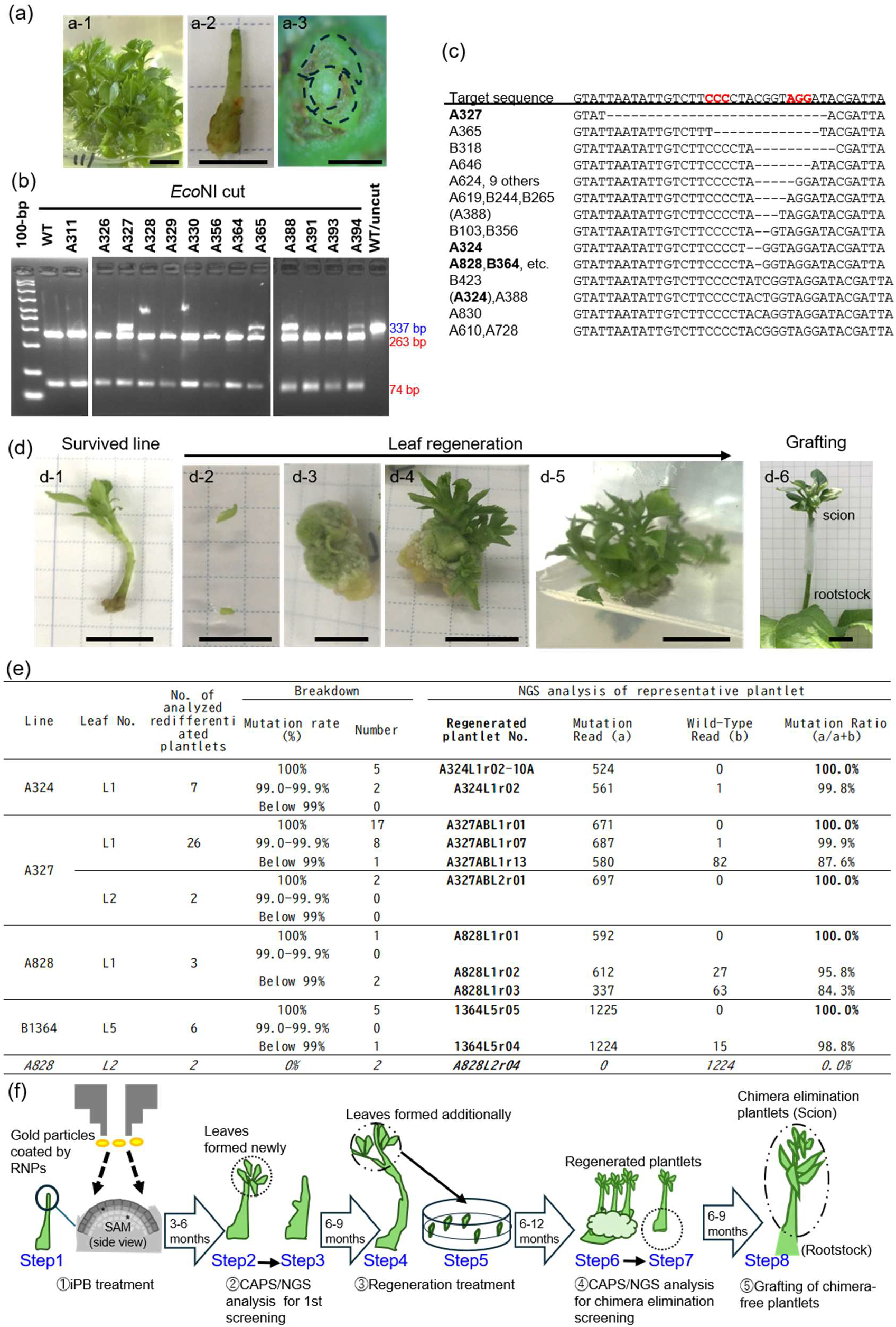
DNA-free iPB-RNP-mediated genome editing of the *S9-RNase* gene in the apple cultivar ‘Fuji’ and regeneration-based elimination of chimerism. (a) Preparation of in vitro-propagated ‘Fuji’ shoots for iPB-RNP treatment. (a-1) In vitro-propagated shoot. Scale bar = 1 cm. (a-2) Shoot with an exposed shoot apical meristem (SAM). Scale bar = 1 cm. (a-3) Top view of the exposed SAM. Dashed outlines indicate the positions of the excised leaves used to expose the SAM. Scale bar = 1 mm. (b) CAPS screening of newly formed leaves following iPB-RNP bombardment. The sizes of undigested (blue) and EcoNI-digested (red) fragments are indicated. (c) MiSeq-based mutation analysis of CAPS-positive shoots. The wild-type sequence is shown at the top. Minor mutations detected within the same line are indicated in parentheses. Mutation frequencies and sequence information are provided in Tables S1 and S2. (d) Regeneration-mediated dechimerization of edited lines. (d-1) Surviving edited line. (d-2 to d-5) Regeneration process from leaf explants. (d-6) Grafted plant on the ‘JM1’ rootstock. Scale bars = 1 cm. (e) MiSeq analysis of regenerated plantlets. The basal portion of the regenerated plantlets (step 7 in (f)) was excised and subjected to MiSeq analysis. Detailed data are provided in Table S4. (f) Schematic overview of the iPB-REG workflow. The numbers of plants at each step are summarized in Table S5.

A total of 1,662 bombarded shoots were cultured in vitro until at least five leaves had developed. Based on the 2/5 phyllotaxy of apple (Okabe, 2015), five consecutive leaves from each shoot were subjected to mutation analysis. Leaf samples were initially screened using cleaved amplified polymorphic sequence (CAPS) analysis (Fig. 1b), and shoots exhibiting undigested bands were subsequently analyzed by MiSeq amplicon sequencing (Tables S1 and S2). Thirty-four edited shoots were identified and are hereafter designated as edited lines (Fig. 1c). These lines contained 18 distinct mutation types (Table S1), with mutation frequencies reaching up to 24.2% (line A744; Table S2). Four lines harbored two independent mutation events. For example, line A324 contained a 2-bp deletion and a 1-bp insertion at frequencies of 16.1% and 3.3%, respectively.

Because the edited shoots were chimeric, a dechimerization step was required to obtain stable mutant plants. To achieve this, a leaf tissue-based regeneration approach was employed. Of the 34 edited lines, 16 were lost during subsequent culture, whereas the remaining 18 produced between one and nine newly formed leaves suitable for regeneration (Fig. 1d-1, Table S3). In total, 96 leaves from these 18 lines were subjected to regeneration (Fig. 1d-2) and cultured for 6-12 months with monthly subculturing (Fig. 1d-3 to d-5). Prior to regeneration, a portion of tissue from each of 84 leaf explants was excised for CAPS analysis. Mutations were detected in 13 leaves, two of which subsequently regenerated into plants (A327L1 and B364L5) (Table S3). To avoid potential reductions in shoot regeneration capacity through tissue excision, the remaining 12 leaves were transferred directly to regeneration medium without prior sampling for CAPS analysis. Eleven of these 12 leaves regenerated successfully, corresponding to a regeneration rate of 92% (11/12). Regenerated shoots were subsequently analyzed by amplicon sequencing.

MiSeq analysis was performed to evaluate the chimeric status of the regenerated plants (Table S4). In addition to lines A327 and B364, regenerated plantlets derived from lines A324 and A828 contained no detectable wild-type reads among more than 300 sequencing reads, indicating the apparent elimination of chimerism (Fig. 1e, Table S4). Notably, regeneration from two independent leaves of line A828 produced either a chimera-free edited plantlet (A828L1r01) or a wild-type plantlet (A828L2r04), demonstrating the segregation of edited and non-edited cell populations during regeneration. No direct correlation was apparent between the mutation frequency in the original chimeric shoot and the successful recovery of dechimerized plants. Representative dechimerized plantlets were grafted for future phenotypic evaluation of self-compatibility (Fig. 1d-6).

By combining the direct delivery of CRISPR-Cas9 RNPs into shoot apical meristems with regeneration-mediated dechimerization and NGS-assisted selection, four complete *S9-RNase*-edited lines were successfully recovered from 1,662 bombarded SAMs (Table S5) in the elite apple cultivar ‘Fuji’. These results demonstrate that iPB-REG can overcome the challenge of chimerism in RNP-based DNA-free genome editing, providing a practical route for the precise genetic improvement of clonally propagated fruit trees while preserving elite cultivar identity. The complete workflow for this strategy is summarized in Fig. 1f.

## Acknowledgements

The authors thank Dr. Sadao Komori for his valuable advice regarding the culture and regeneration of the apple cultivar ‘Fuji’. Recombinant SpCas9 protein was generously provided through the Advanced Analysis Center Research Support Program of the National Agriculture and Food Research Organization (NARO).

## Author Contributions

C.N. and R.I. conceived and designed the study. C.N., N.T., M.K., and M.W. performed the experiments. C.N. analyzed the data. C.N. and R.I. wrote the manuscript.

## Supporting Information

### Materials and Methods

#### Preparation of Apple Shoot Apical Meristems (SAMs)

Shoot apical meristems (SAMs) were prepared from in vitro-propagated apple (*Malus × domestica*) ‘Fuji’ plantlets. To expose the SAMs, several small leaves covering the apex were carefully removed under a stereomicroscope using an insulin pen needle (34G; TERUMO, Japan). The shoots were cut into 1-cm segments and placed upright in Petri dishes containing Murashige and Skoog (MS) basal medium supplemented with maltose (30 g/L), 2-(N-morpholino)ethanesulfonic acid (MES) monohydrate (0.98 g/L; pH 5.8), plant preservative mixture (3%; Nacalai Tesque, Japan), and Phytagel (7.0 g/L; Sigma-Aldrich, USA). Approximately twenty shoots were placed in each dish for each round of particle bombardment.

#### Coating of Gold Particles with sgRNA-SpCas9 Complex and Particle Bombardment

Recombinant *Streptococcus pyogenes* Cas9 protein was purified from *Escherichia coli* as previously described (Hamada et al., 2017). CRISPR RNA (crRNA) and trans-activating CRISPR RNA (tracrRNA) were obtained from Integrated DNA Technologies (Japan). To hybridize the RNAs, crRNA and tracrRNA, each at 100 μM, were mixed, heated at 95°C for 5 min, and slowly cooled according to the manufacturer’s protocol. Preparation of the RNP complex and coating to gold particles were performed according to Kumagai et al. (2022) with slight modifications. Particle bombardment was performed using a Biolistic PDS-1000/He system (Bio-Rad). The distance between the plantlets and the brass adjustable nest of the microcarrier launch assembly was approximately 3.5 cm. Each plate was bombarded four times at a helium pressure of 1,300 psi. After bombardment, the explants were transferred to MS medium (pH 5.6) and subcultured monthly.

#### Selection and Evaluation of Genome-Edited Plantlets

Genome-edited plantlets were screened using cleaved amplified polymorphic sequence (CAPS) analysis. Several leaves surrounding each shoot were collected for genomic DNA extraction. The target region of the *S9-RNase* gene was amplified using the following primers: 5′-ATGGGGATTACGGGGATGATATATATGGTTA-3′ and 5′-TTCACTAGCATGCATGAAAATCTATGTTGAGTATATG-3′. For gel-based CAPS analysis, the primers were used without modification, whereas for analysis using an ABI 3130xl Genetic Analyzer, the forward primer was labeled with a fluorescent dye (FAM). PCR was performed in a 10-μL reaction volume using PrimeSTAR GXL DNA Polymerase (Takara Bio). The first PCR consisted of an initial denaturation at 95°C for 3 min, followed by 25 cycles of 95°C for 30 s, 58°C for 30 s, and 72°C for 30 s, with a final extension at 72°C for 5 min. One microliter of the first PCR product was used as the template for the second PCR, which was performed under identical conditions. The amplified DNA fragments were digested with EcoNI (New England Biolabs) at 37°C for 16 h, followed by enzyme inactivation at 65°C for 20 min. The digested products were analyzed either by 2% agarose gel electrophoresis with GelRed staining (Biotium) or using an ABI 3130xl Genetic Analyzer.

For detailed mutation analysis, amplicon sequencing was performed using the MiSeq platform (Hokkaido System Science). Libraries were prepared according to the manufacturer’s instructions. Indexed PCR products were pooled, sequenced, and assigned to individual samples based on their index sequences. Reads containing both primer sequences were retained for analysis. Because MiSeq data processing treats even single-nucleotide differences outside the target site as distinct sequences, sequence variants unrelated to the target site and likely attributable to PCR or sequencing errors were excluded. The region corresponding to the 38-bp wild-type sequence containing the target site was then trimmed and sequence counts were summarized (Hokkaido System Science). Sequence variants within this region were classified according to the mutation types defined in Table S1, and reads that could not be assigned to any defined mutation type were classified as “Unclassified” in Tables S2 and S4. The sequencing data have been deposited in the NCBI database (accession numbers: DRR913357–DRR913444).

In the CAPS/NGS analysis used for the first screening (Fig. 1c, 1g; Table S2), variants representing ≥4% of the total reads in each sample were counted as mutation events, because this threshold empirically corresponded to unequivocal CAPS-positive signals rather than incomplete restriction digestion. For chimera-elimination screening, regenerated plantlets were first examined by CAPS analysis, and CAPS-positive candidates were subsequently analyzed by next-generation sequencing (NGS). Based on a previous amplicon-sequencing study of chimerism in genome-edited transgenic apple, in which mutation frequencies were evaluated at approximately 1% resolution (Pompili et al., 2019), only samples with sufficient sequencing depth to reliably detect low-frequency variants were included. Accordingly, further analysis was limited to only samples containing ≥300 total reads and no detectable wild-type reads (Fig. 1e, g; Table S4).

#### Redifferentiation of Leaves Derived from Mutation-Retaining Shoots

Leaves collected from shoots confirmed to retain genome-edited mutations were subjected to redifferentiation according to the protocol described by Li et al. (2023). Briefly, leaf segments were cultured on redifferentiation medium in the dark for 2 weeks. The cultures were then transferred to low-light conditions under a 16-h light/8-h dark photoperiod for an additional 2 weeks. Subsequently, the leaf segments were transferred to shoot regeneration medium and subcultured monthly under the same photoperiod.

**Supplementary Figure 1.**
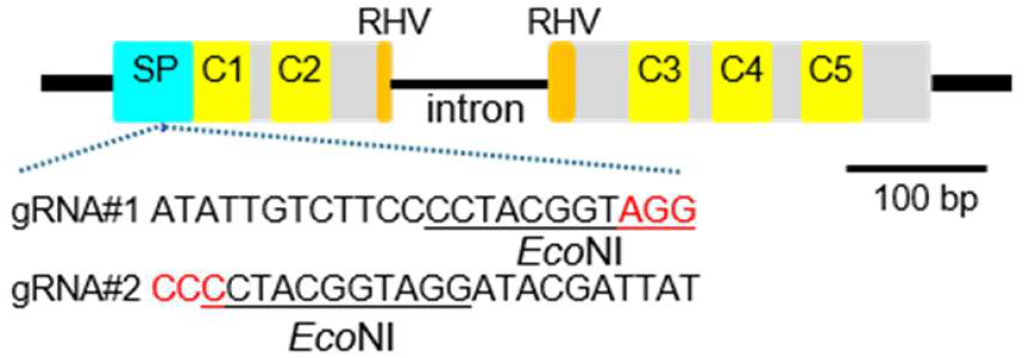
Structure of the *S9-RNase* gene showing the target sites of gRNA#1 and gRNA#2. Red letters indicate the Cas9 PAM (NGG), and the EcoNI recognition site used for CAPS analysis is underlined. SP, putative signal peptide region; C1–C5, conserved regions; RHV, hypervariable region.

## Supplemental Tables

**Table S1.**
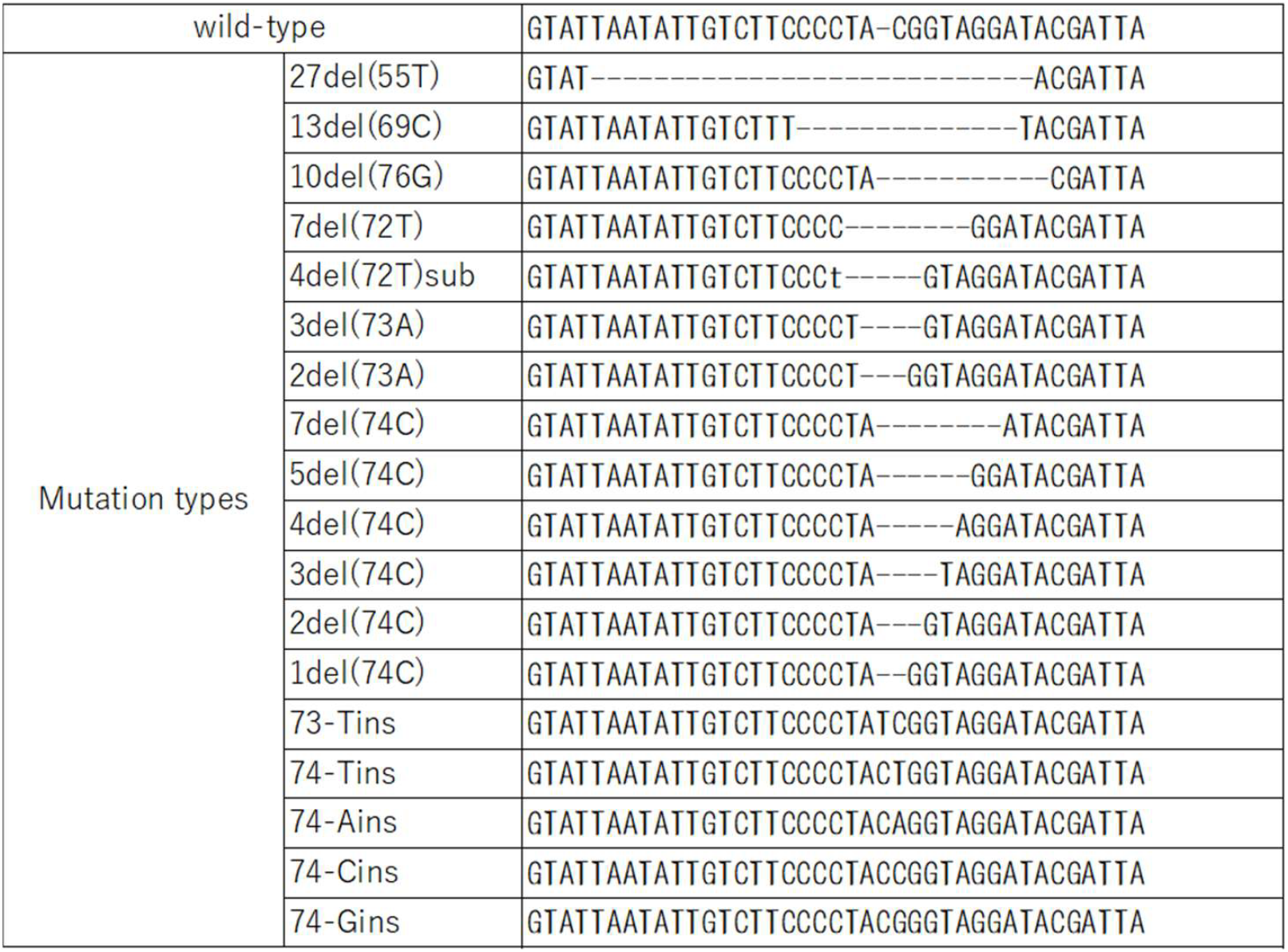
Types of detected mutations.

**Table S2.**
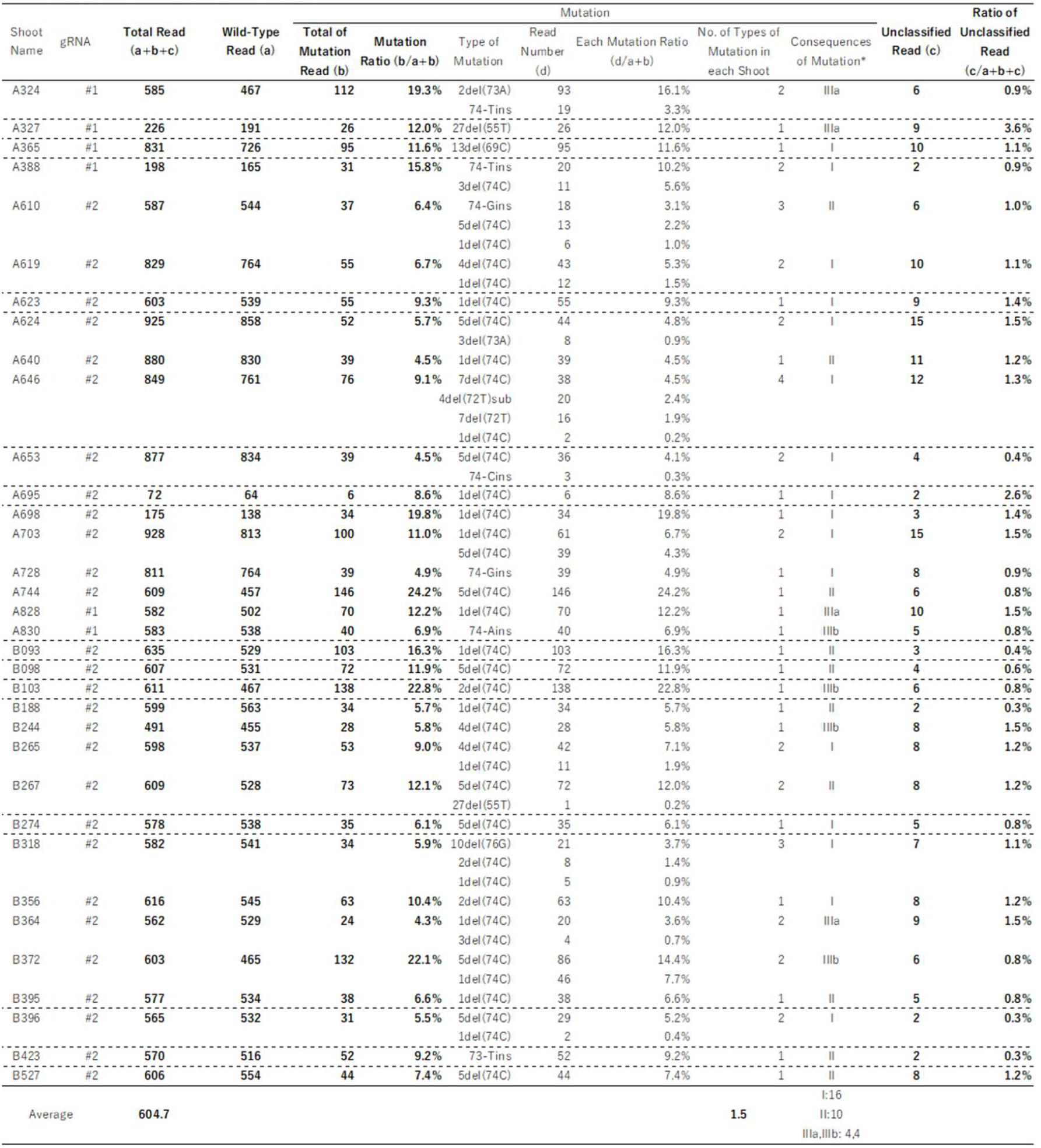
Amplicon sequence analysis of bombarded plantlets.

**Table S3.** Number of regenerated shoots.

| Line | CAPS | CAPS positive |  | CAPS negative |  | Not analyzed |  |
| --- | --- | --- | --- | --- | --- | --- | --- |
|  | Regeneration | + | - | + | - | + | - |
| A324 |  | **0** | 1 | **0** | 2 | **1(L1)** | 0 |
| A327 |  | **1(L1)** | 1 | **0** | 6 | **1(L2)** | 0 |
| A610 |  | **0** | 0 | **1** | 0 | **0** | 0 |
| A640 |  | **0** | 0 | **0** | 6 | **1** | 0 |
| A744 |  | **0** | 0 | **0** | 4 | **1** | 0 |
| A828 |  | **0** | 2 | **0** | 1 | **2(L1, L2)** | 1 |
| A830 |  | **0** | 2 | **0** | 5 | **0** | 0 |
| B093 |  | **0** | 0 | **0** | 4 | **1** | 0 |
| B098 |  | **0** | 0 | **0** | 1 | **1** | 0 |
| B103 |  | **0** | 1 | **0** | 0 | **0** | 0 |
| B188 |  | **0** | 0 | **0** | 3 | **0** | 0 |
| B244 |  | **0** | 2 | **0** | 5 | **0** | 0 |
| B267 |  | **0** | 0 | **0** | 0 | **3** | 0 |
| B364 |  | **1(L5)** | 1 | **0** | 7 | **0** | 0 |
| B372 |  | **0** | 1 | **0** | 4 | **0** | 0 |
| B395 |  | **0** | 0 | **0** | 8 | **0** | 0 |
| B423 |  | **0** | 0 | **0** | 9 | **0** | 0 |
| B527 |  | **0** | 0 | **0** | 5 | **0** | 0 |
| Total |  | **2** | 11 | **1** | 70 | **11** | 1 |

**Table S4.**
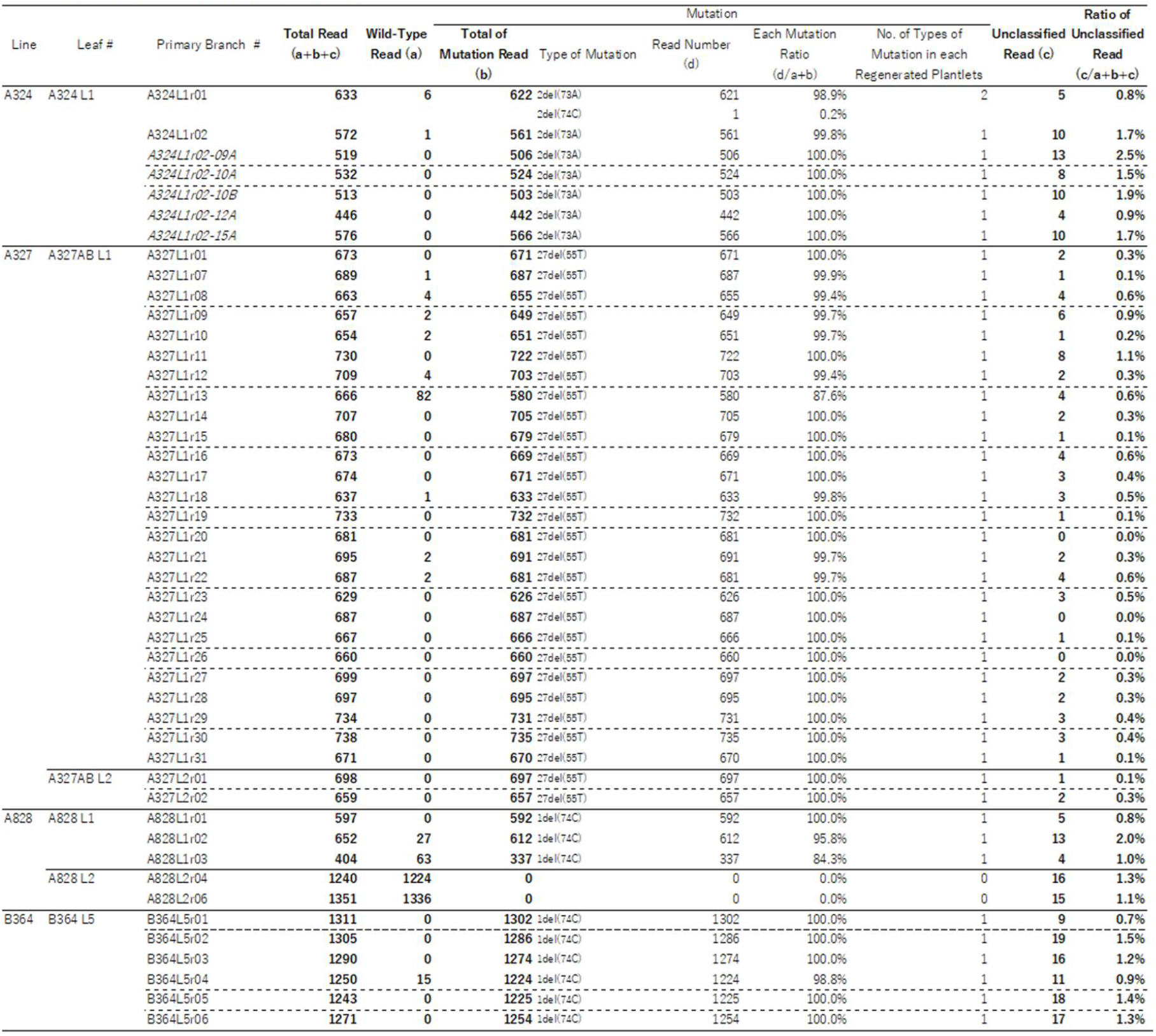
Amplicon sequence analysis of regenerated plantlets.

**Table S5.** Summary of iPB-RNP approach to S9-RNase editing in ‘Fuji’.

| Steps | No. of candidate plant |
| --- | --- |
| 1 Initially bombarded SAMs | 1,662 |
| 2 Shoots available for CAPS analysis | 1,162 |
| 3 CAPS-positive shoots | 34 |
| 4 Survived and subjected to leaf regeneration | 18 |
| 5 Regeneration from leaf explants | 10 |
| 6 Regenerated plantlets analyzed: 91 (from 10 lines) | 10 |
| 7 NGS-confirmed chimera-free plantlets (f): 30 (from 4 lines) | 4 |
| Total efficiency (%) | 0.24 |

## References

Broothaerts, W., Keulemans, J., & Van Nerum, I. (2004). Self-fertile apple resulting from S-RNase gene silencing. Plant Cell Reports, 22, 497–501.

Kumagai et al. (2022).Introduction of a second “Green Revolution” mutation into wheat via in planta CRISPR/Cas9 delivery. Plant Physiology, 188, 1838–1842.

Kuwabara, C., Miki, R., Maruyama, N., Yasui, M., Hamada, H., Nagira, Y., Li, F., Hirayama, Y., Ackley, W., Feng, L., & Imai, R. (2024). DNA-free and genotype-independent CRISPR/Cas9 system in soybean. Plant Physiology, 196(4), 2320–2329.

Imai, R., Hamada, H., Liu, Y., Linghu, Q., Kumagai, Y., Nagira, Y., Miki, R., & Taoka, N. (2020). In planta particle bombardment (iPB): A new method for plant transformation and genome editing. Plant Biotechnology, 37, 171–176.

Malabarba, J., Chevreau, E., Dousset, N., Veillet, F., Moizan, J., & Vergne, E. (2021). New strategies to overcome present CRISPR/Cas9 limitations in apple and pear: efficient dechimerization and base editing. International Journal of Molecular Sciences, 22(1), 319.

Nishitani, C., Hirai, N., Komori, S., Wada, M., Okada, K., Osakabe, K., Yamamoto, T., & Osakabe,Y. (2016). Efficient genome editing in apple using a CRISPR/Cas9 system. Scientific Reports, 6, 31481.

Okabe, T. (2015). Biophysical optimality of the golden angle in phyllotaxis. Scientific Reports, 5, 15358. 10.1038/srep15358

Okada, K., Shimizu, T., Moriya, S., Wada, M., Abe, K., & Sawamura, Y. (2024). Alternative splicing and deletion in S-RNase confer stylar-part self-compatibility in the apple cultivar ‘Vered’. Plant Molecular Biology, 114, Article 113.

Pompili, V., Dalla Costa, L., Piazza, S., Pindo, M., & Malnoy, M. (2019). Reduced fire blight susceptibility in apple cultivars using a high-efficiency CRISPR/Cas9-FLP/FRT-based gene editing system. Plant Biotechnology Journal, 18, 845–858. 10.1111/pbi.13253

Sasaki, K., Urano, K., Mimida, N., Nonaka, S., Ezura, H., & Imai, R. (2025). A long shelf-life melon created via CRISPR/Cas9 RNP-based in planta genome editing. Frontiers in Genome Editing, 7, 1623097.

## Supplemental References

Hamada, H., Linghu, Q., Nagira, Y., et al. (2017). An in planta biolistic method for stable wheat transformation. Scientific Reports, 7: 11443. 10.1038/s41598-017-10930-5

Li, F., Kawato, N., Sato, H., Kawaharada, Y., Henmi, M., Shinoda, A., … & Komori, S. (2023). Release of chimeras and efficient selection of editing mutants by CRISPR/Cas9-mediated gene editing in apple.Scientia Horticulturae, 316, 112011.

